# Chemical structure elucidation of the carotenoids in the extremely radiation-resistant bacterium *Deinococcus radiodurans* R1

**DOI:** 10.1101/2021.05.26.445707

**Authors:** Xiao-ling Zhang, Qiao Yang

**Author notes:** Corresponding author. E-mail addresses (Q. Yang).

## Abstract

The main carotenoid in the extremely radiation-resistant bacterium *Deinococcus radiodurans* is deinoxanthin. In this paper, based on UV-Vis, MS/MS spectra as well as the chemical properties, the other main two carotenoids in this bacterium were identified as ((all-E)-1’-hydroxy-3’,4’-didehydro-1’,2’-dihydro-β,Ψ-carotene- 4-one-1), and ((all-E)-1’-hydroxy-3’,4’-didehydro-1’,2’,2,3-quahydro-β,Ψ-carotene- 4-one-1), respectively. The fully structure elucidation for the carotenoids should help to gain a better understanding of the radiation-resistant mechanisms in this remarkable bacterium.

## Introduction

*Deinococcus radiodurans* is a gram-positive red-pigmented, nonspordforming, nonmotile spherical bacterium that was originally identified as a contaminant of canned meat irradiated with a sterilizing dose of γ radiation. *D. radiodurans* is a polyextremophile, showing remarkable resistance to a range of several damage caused by ionizing radiation, desiccation, UV radiation, a number of agents and conditions that damage DNA, or electrophilic mutagens. This bacterium is the most radiation-resistant organism described to data; exponentially growing cells are 200 times as resistant to ionizing radiation and 20 times as resistant to UV irradiation as *Escherichia coli*.

Carotenoids represent one of the most widely distributed and structurally diverse classes of natural pigments, producing colors ranging from light yellow through orange to deep red. At least 600 different carotenoids are synthesized naturally, performing important functions in photosynthesis, nutrition and protecting against oxidative damage. The instability of the carotenoids to heat, light and air make isolation, identification and quantitation of these natural pigments unusually challenging. Many carotenoids exhibit provitamin A activity, others might function as significant dietary antioxidants, and other beneficial effects of carotenoids that are currently under investigation include inhibition of carcinogenesis, enhancement of the immune response and prevention of cardiovascular disease. Recent transcriptome analysis revealed that *Deinococcus radiodurans* efficiently coordinate their recovery from ionizing radiation through a complex network of DNA repair and metabolic pathway switching. However, the additional discovery of numerous irradiation response genes has provided new targets for the identification of genes primarily crucial to radiation resistance. Although such investigations provide useful insights into the mechanisms underlying radiation resistance, a more detailed empirical explanation of why *D. radiodurans* is so radiation resistant is still needed.

Further research based on alternative genetic and biochemical approaches should help to gain a better understanding of the mechanisms involved in DNA repair The main carotenoids pigment of *Deinococcus radiodurans* has been identified as deinoxanthin ((all-E)-2,1’-dihydroxy-3’,4’-didehydro-1’,2’-dihydro-β,Ψ-carotene-4-one-1) by Lemee et al. however, the chemical structures of other carotenoids in this bacterium have not been reported, The present work is to deal with the determination of the chemical structures of the red pigments in *D. radiodurans*.

## Materials and methods

### Chemicals and reagents

Acetonitrile, dichloromethane and methanol (HPLC grade) was obtained from Fisher (Fair Lawn, NJ, USA). Acetone was purchased from Merck (Darmstadt, Germany). Deionized water from a Milli-Q system (Millipore, Bedford, MA, USA) was used to prepare all buffers and samples solutions.

### Bacteria strain and growth conditions

The R1 strain of *Deinococcus radiodurans* was obtained from CCTCC (China Center for Type Culture Collection, Wuhan, China) was grown in TGY media (0.5% tryptone, 0.3% yeast extract, 0.1% glucose, pH 7.2) with aeration on rotary shakers or on TGY plates (TGY broth supplemented with 1.5% agar) at 30 °C.

### Extraction of carotenoids from *D. radiodurans*

After 48 hours of incubation at 30 °C under aerobic conditions and shaking (250 rpm), the culture was harvested by centrifugation (4000 rpm, 20 min), after washing three times with deionized and sterilized water, the cell pellet was mixed with a 20 ml mixture of cold acetone and methanol (3:2, v/v) with 0.1% BHT(buthlated hydroxytoluene) as antioxidant, blended in a mechanical blender for 20 minutes, and then centrifuged (6000 rpm, 20 min at 4 °C). The extraction was repeated for three times until the residue was devoid of any color and washing were colorless. The extracts were combined and concentrated under vacuum in a rotary evaporator to dryness in the dark at 30 °C. The extract was transferred quantitatively to a 10 ml volumetric flask using portions of 2 ml of mobile phase for the sequent analysis.

### HPLC and LC/MS/MS analysis of carotenoids in *D. radiodurans*

The HPLC analysis was performed on an Agilent 1100 high-performance liquid chromatograph (Agilent, Palo Alto, CA, USA) equipped with a diode-assay detector (DAD) covering the range of 190–600 nm. Separation was carried out on an Agilent reverse-phased ODS C_18_ column (5 μm, 4.6 mm × 250 mm I.D.) maintained at 25 °C. A gradient consisting of solvent A: 85% acetonitrile, 10% dichloromethane and 5% methanol; solvent B: water, 0-12 min 100% A; 12-35 min 85% A was used. The mobile phase consisting of acetonitrile-dichloromethane-methanol (85:10:5, v/v/v) was used for the isocratic elution. The flow rate was 1.0 mL min^-1^and the UV-visible DAD detection was monitored at 470 nm.

The semipreparative separation of the carotenoids was performed on Waters Delta 600E semi-HPLC equipped with PDA detector, an autosampler, and a fraction collector. A Symmetry Prep™ reverse-phased ODS C_18_column (7 μm, 7.8 mm × 300 mm I.D.) was used. The Empower Pro was used for the instrument control and data analysis. The purified major product was identified by reversed phase HPLC with a purity of 97%.

The LC/MS/MS analysis of carotenoids was performed using a Finnigan LCQ™ Deca XP MAX ion trap mass spectrometer (Thermo Finnigan, CA, USA) equipped with an atmospheric pressure chemical ionization (APCI) interface (Finnigan), which connected with a Finnigan Surveyor HPLC system equipped with a quaternary pump, a vacuum membrane degasser, a LightPipe™ UV-visible photodiode-array (PDA) detector, an autosampler. Chromatography was carried out on a column with a 2.1 mm ×5.0 mm I.D. guard column. A gradient consisting of solvent A: 85% acetonitrile, 10% dichloromethane and 5% methanol; solvent B: water, 0-12 min 100% A; 12-35 min 85% A was used. Multiple reaction monitoring (MRM) was used to perform mass spectrometric identification of a unique product ion from a selected molecular ion that can be monitored and quantified even in the midst of a very complicated biological matrix. The column effluent was introduced into the mass spectrometer using an APCI interface in the positive mode. The voltage of source was 6000 V and the capillary voltage was 19 V. Selected [M+H]^+^ was analyzed by collision-induced dissociation with argon gas, and product ion spectra were recorded.

## Results and discussion

### HPLC analysis of carotenoids of *D. radiodurans*

The carotenoids isolated from the cell culture of *D. radiodurans* was separated and detected by HPLC with diode array detection at 470 nm. Three main peaks were found in the HPLC chromatogram (Fig. 1). The UV/Vis spectra of peak 1, 3 and 4 were compared with that of deinoxanthin (peak 2), we found the maximum of peak 1 and 4 are almost the same as that of deinoxanthin, indicating that they have very close chemical structure. That is, they may have the same number of the conjugated polyene system. Peak 2 is the cis isomer of deinoxanthin, as they have the same molecular ion M+ at *m/z* at 582 and the maxima at 451, 580 and 506 nm from the PDA detection, whereas peak 3 have another UV/Vis spectrum at 270 nm that is the characteristic spectra of the cis isomer of a carotenoids.

**Figure 1.**
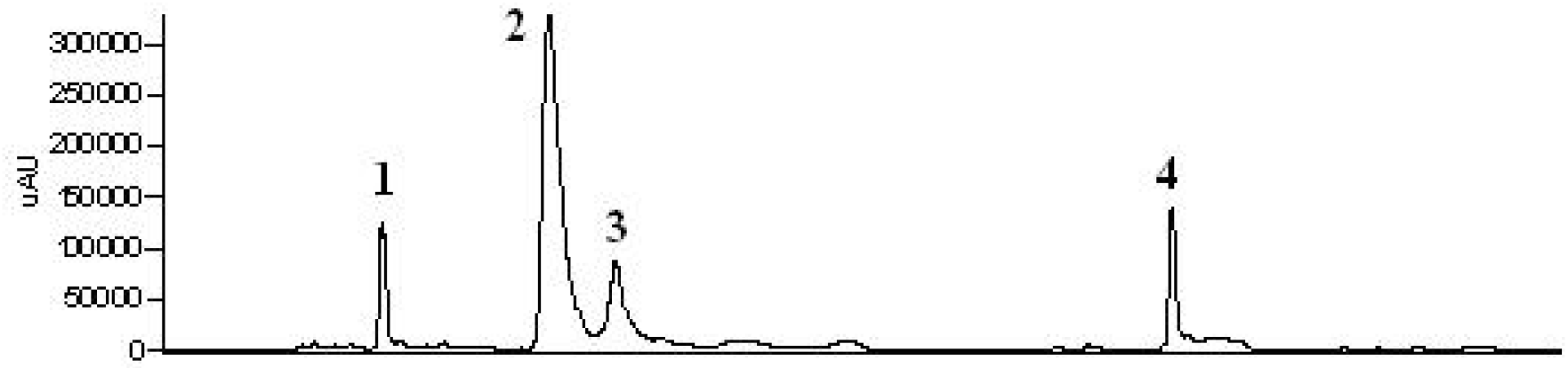
HPLC chromatogram of carotenoids of *D. radiodurans*. Conditions were as described in the “Materials and methods’ section.

### HPLC-tandem MS analysis of carotenoids in *D. radiodurans*

Carotenoids of *D. radiodurans* were analyzed with both positive and negative ion modes. They all showed the protonated molecule [M+H]^+^ in the positive ion mode. In the negative ion mode, their deprotonated molecules were very weak. Therefore, the positive ion mode was utilized for the carotenoids analysis.

In the MS/MS experiment, For peak 1, the *m/z* at 565.3 ([M+H]^+^) corresponding to a molecular formula C_40_H_52_O_2_ produced a cluster of ions at *m/z* 546, 505, 401 and along with a series of product ions in the mass range of *m/z* 100-300. Further characteristic fragmentation were observed at *m/z* 547.2 ([M+H]^+^-18) indicating the loss of H_2_O. For peak 3, the *m/z* at 583.2 ([M+H]^+^) corresponding to a molecular formula C_40_H_54_O_3_. Further characteristic fragmentation were observed at *m/z* 547.2 ([M+H]^+^-18) indicating the loss of H_2_O. For peak 4, the *m/z* at 567.3 ([M+H]^+^) corresponding to a molecular formula C_40_H_54_O_2_. Further characteristic fragmentation were observed at *m/z* 548.3 ([M+H]^+^-18) indicating the loss of H_2_O. Based on the UV-Vis, and MS/MS spectra as well as the chemical properties the two carotenoids were identified as, ((all-E)-1’-hydroxy-3’,4’-didehydro-1’,2’-dihydro-β,Ψ-carotene-4-one-1) for peak 1, ((all-E)-1’-hydroxy-3’,4’-didehydro-1’,2’,2,3-quahydro-β,Ψ-carotene-4-one-1) for peak 4, respectively (Fig. 4). The proposed biosynthesis pathway was also shown.

**Figure 2.**
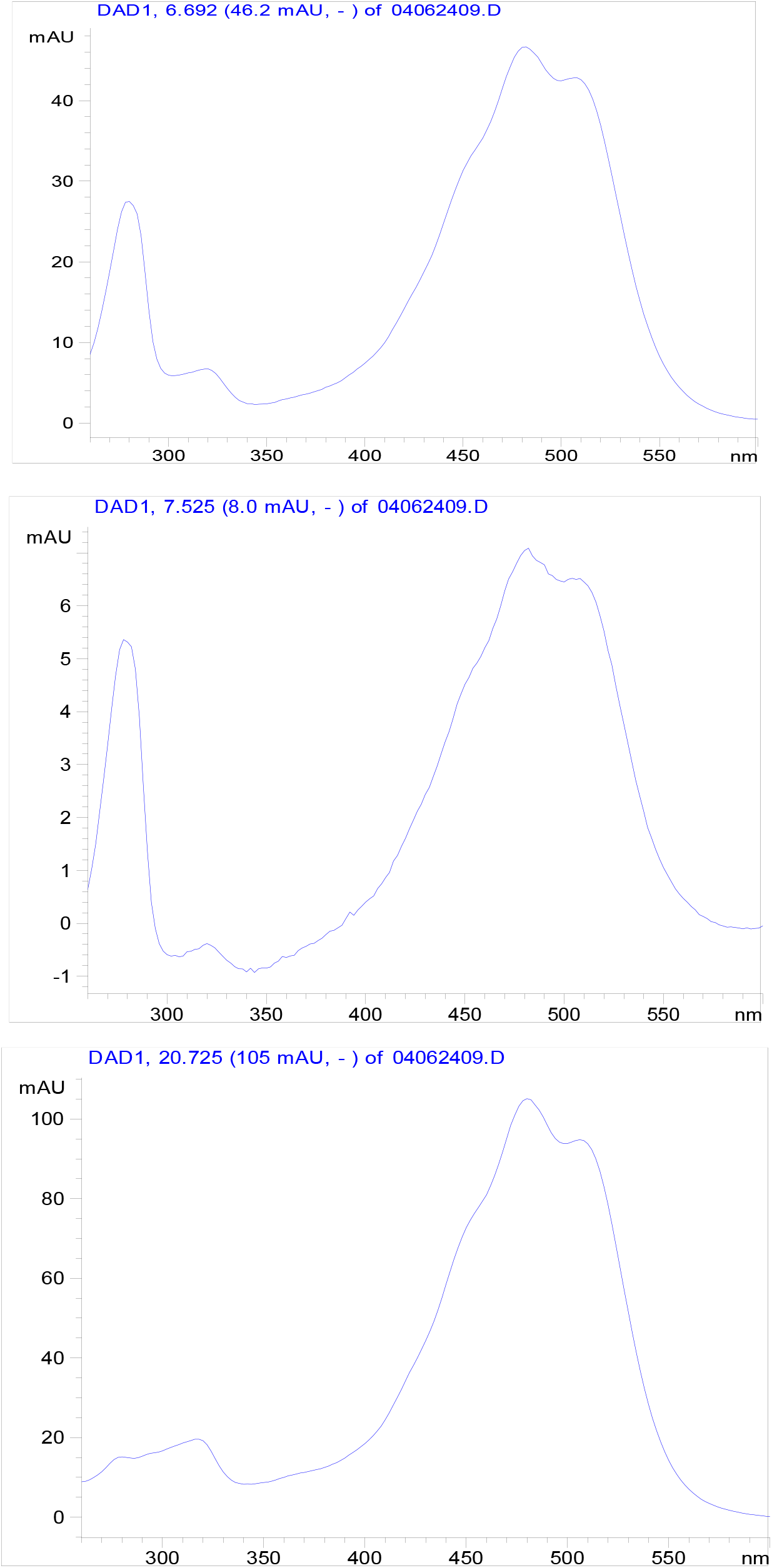
UV-Vis spectra of four peaks from the HPLC chromatogram of carotenoids in *D. radiodurans*.

**Figure 3.**
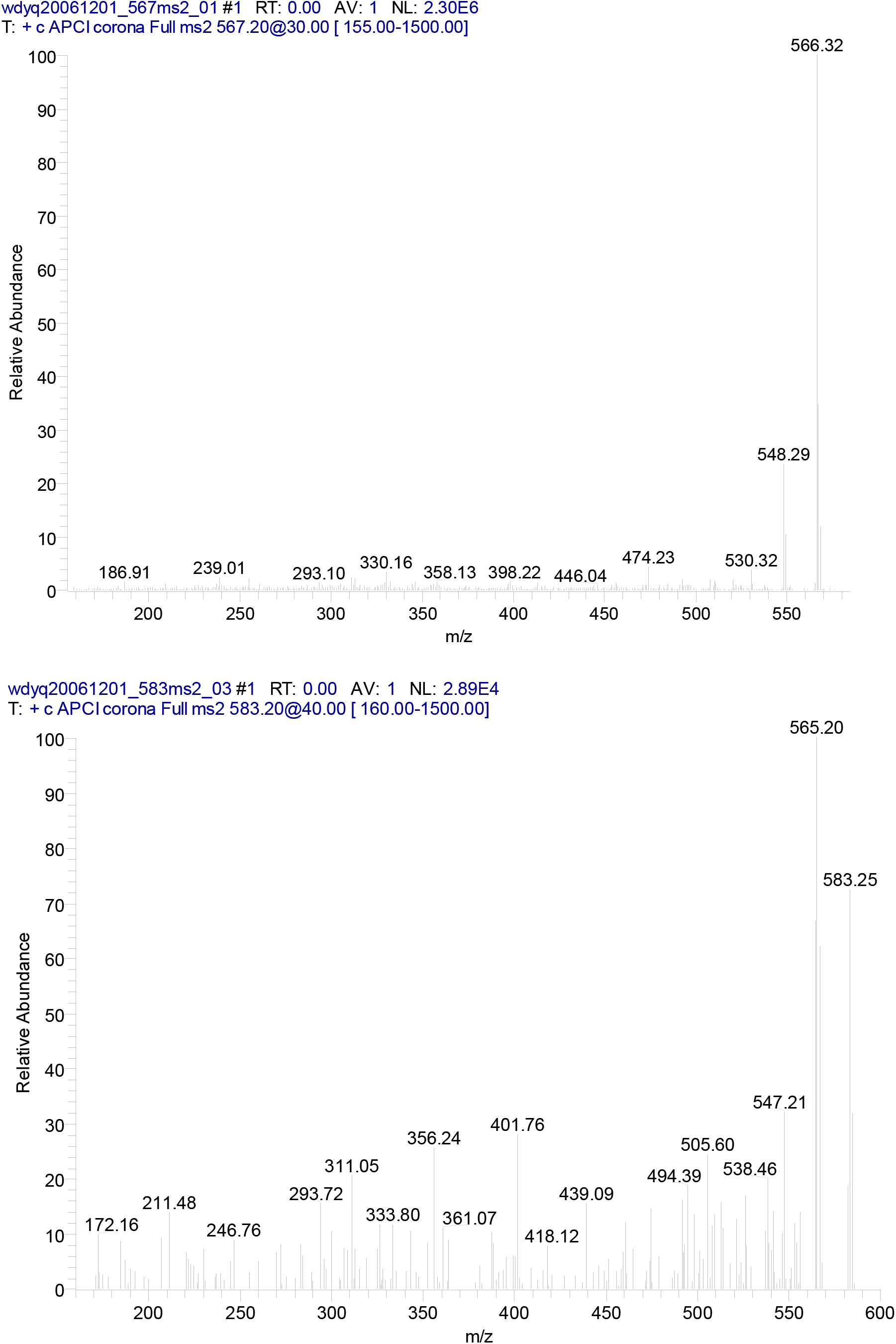
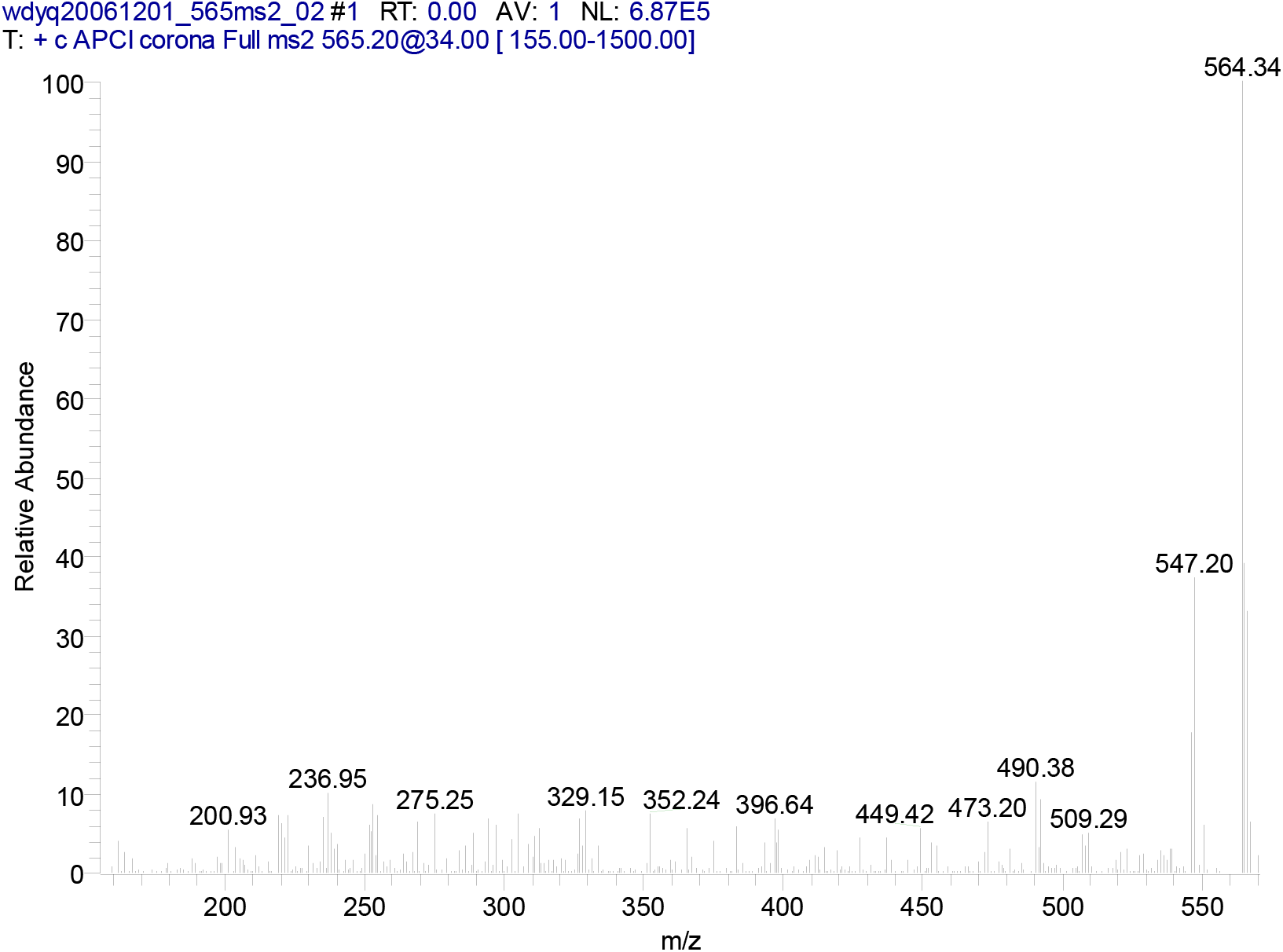
Mass spectra of the carotenoids in *D. radiodurans*.

**Figure 4.**
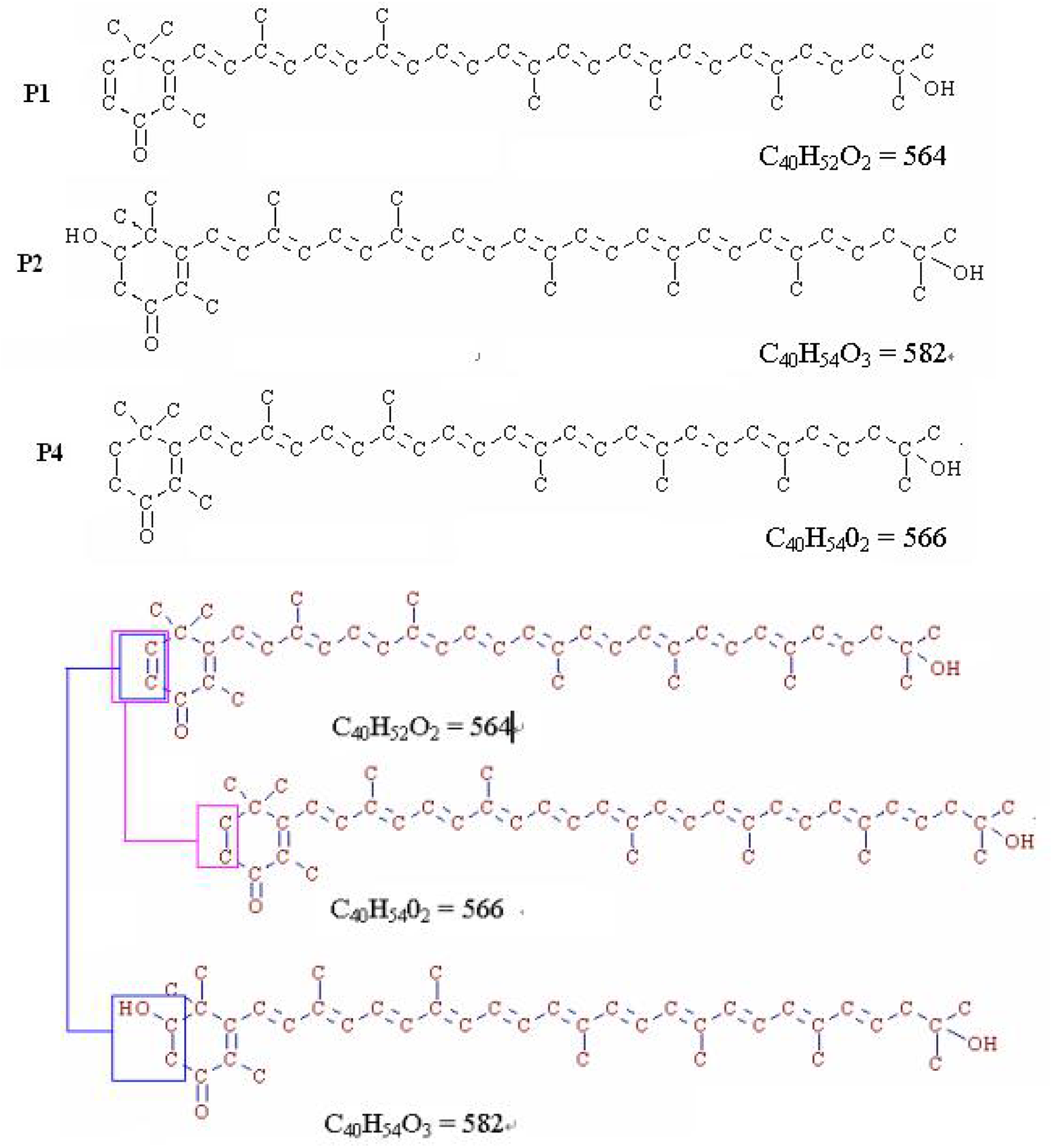
The chemical structures of the main carotenoids in *D. radiodurans* and the proposed biosynthesis pathway.

## Conclusion

It has been shown that carotenoids appear, at least in part, as a radioprotective agent. Ionizing radiation can induce singlet oxygen and free radicals and carotenoids were initially known as a photo reductone and a singlet oxygen quencher. It has been widely accepted that very strong relationships exist between the chemical structure and antioxidant function of carotenoids. The two main carotenoids besides deinoxanthin were isolated and analyzed by UV-Vis, APCI-MS/MS as well as the chemical properties test to obtain structure information. This study will help to gain a better understanding of the radiation-resistant mechanisms in this remarkable bacterium.

## References

1 Anderson A W, Nordan H C, Cain R F, et al. Studies on a radio-resistant micrococcus: isolation, morphology, cultural characteristics, and resistance to gamma radiation. Food Technol, 1956, 10: 575∼578

2 Battista J R. Against all odds: the survival strategies of Deinococcus radiodurans. Annu Rev Microbiol, 1997, 51: 203∼220

3 Cox M M, Battista J R, Deinococcus radiodurans - the consummate survivor. Nat Rev Microbiol, 2005, 3(11): 882∼892

4 White O, Eisen J A, Heidelberg J F, et al. Genome sequence of the radioresistant bacterium Deinococcus radiodurans R1. Science, 1999, 286(5444): 1570∼1577

5 Lange C C, Wackett L P, Minton K W, et al. Engineering a recombinant Deinococcus radiodurans for organpollutant degradation in radioactive mixed waste environments. Nat Biotechnol, 1998, 16(10): 929∼933

6 Brim H, McFarlan S C, Fredrickson J K, et al. Engineering Deinococcus radiodurans for metal remdiation in radioactive mixed waste environments. Nat Biotechnol, 2000, 18(1): 85∼90

7 Ghosal D, Omelchenko M V, Gaidamakova E K, et al. How radiation kills cells: Survival of Deinococcus radiodurans and Shewanella oneidensis under oxidative stress. FEMS Microbiol Rev, 2005, 29(2): 361∼375

8 Kolari M, Schmidt U, Kuismanen E, et al. Firm but slippery attachment of Deinococcus geothermalis. J Bacteriol, 2002, 184(9): 2473∼2480

9 Funayama T, Narumi I, Kikuchi M, et al. Identification and disruption analysis of the recN gene in the extremely radioresistant bacterium Deinococcus radiodurans. Mutat Res, 1999, 435(2): 151∼161

10 Carbonneau M A, Melin A M, Perromat A, et al. The action of free radicals on Deinococcus radiodurans carotenoids. Arch Biochem Biophys, 1989, 275(1): 244∼251.

11 Markillie L M, Varnum S M, Hradecky P, et al. Targeted mutagenesis by duplication insertion in the radioresistant bacterium Deinococcus radiodurans: radiation sensitivities of catalase (katA) and superoxide dismutase (sodA) mutants. J Bacteriol, 1999, 181(2): 666∼669

12 Yang Q, Zhang X L, Zhang J X, et al. A novel method for evaluating free radical scavenging abilities of antioxidants using ultraviolet induction of bacteriophage λ. J Biochem Biophys Methods, 2006, 67(2-3): 163∼171

13 Lemee L, Peuchant E, Clerc M, et al. Deinoxanthin: a new carotenoids isolated from Deinococcus radiodurans. Tetrahedrom, 1997, 53(3): 919∼926

14 Yang Q, Zhang J X, Zhu S Q, et al. Study on the radiation resistant substance of Deinococcus radiodurans. J Anal Sci, 2004, 20 (5): 465∼467

15 Umeno D, Tobias A V, Arnold F H. Diversifying carotenoid biosynthetic pathways by directed evolution. Microbiol Mol Biol Rev, 2005, 69(1): 51∼78

16 Makarova K, Aravind L, Wolf Y, et al. Genome of the extremely radiation-resistant bacterium Deinococcus radiodurans viewed from the perspective of comparative genomics. Microbiol Mol Biol Rev, 2001, 65(1): 44∼79

17 Trevithick-Sutton C C, Foote C S, Collins M, et al. The retinal carotenoids zeaxanthin and lutein scavenge superoxide and hydroxyl radicals: a chemiluminescence and ESR study. Mol Vis, 2005, 12: 1127∼1135

18 Tatsuzawa H, Maruyama T, Misawa N, et al. Quenching of singlet oxygen by carotenoids produced in Escherichia col-attenuation of singlet oxygen-mediated bacterial killing by carotenoids. FEBS Lett, 2004, 484(3): 280∼284

19 Moan J. Detection of singlet oxygen production by ESR. Nature, 1979, 279(5712): 450∼451

